# Buffering of transcription rate by mRNA half-life is a conserved feature of Rett syndrome models

**DOI:** 10.1101/2021.12.11.472181

**Authors:** Deivid C. Rodrigues, Marat Mufteev, Kyoko E. Yuki, Ashrut Narula, Wei Wei, Alina Piekna, Jiajie Liu, Peter Pasceri, Olivia S. Rissland, Michael D. Wilson, James Ellis

**Author notes:** equal contribution (Deivid C. Rodrigues, Marat Mufteev).

## Abstract

Models of MECP2 dysfunction in Rett syndrome (RTT) assume that transcription rate changes directly correlate with altered steady-state mRNA levels. However, limited evidence suggests that transcription rate changes are buffered by poorly understood compensatory post-transcriptional mechanisms. Here we measure transcription rate and mRNA half-life changes in RTT patient neurons using RATE-seq, and reinterpret nuclear and whole-cell RNAseq from *Mecp2* mice. Genes are dysregulated by changing transcription rate only or half-life only and are buffered when both are changed. We utilized classifier models to understand the direction of transcription rate changes based on gene-body DNA sequence, and combined frequencies of three dinucleotides were better predictors than contributions by CA and CG. MicroRNA and RNA-Binding Protein (RBP) motifs were enriched in 3’UTRs of genes with half-life changes. Motifs for nuclear localized RBPs were enriched on buffered genes with increased transcription rate. Our findings identify post-transcriptional mechanisms in humans and mice that alter half-life only or buffer transcription rate changes when a transcriptional modulator gene is mutated in a neurodevelopmental disorder.

## Introduction

Rett syndrome (RTT) is a neurodevelopmental disorder in girls caused by damaging mutations in the methyl CpG-binding protein 2 (*MECP2*)^1^. Most evidence indicates that MECP2 regulates transcription globally after binding to methylated (m)CG dinucleotides in immature neurons and mCA dinucleotides in adult neurons^2–8^. Specifically, recent models support a role for MECP2 binding to mCA/mCGs in the gene-body that can: 1) slow RNA polymerase II elongation^5^ 2) inhibit transcription initiation by looping interactions between the promoter and high-density MECP2-bound gene-bodies^9^ or 3) inhibit transcription initiation by microsatellite interruptions of nucleosome binding that impact intragenic enhancer activation^10^.

The number or fraction of mCA/mCG and gene-length have been primarily used to interpret transcription rate dysregulation in RTT mouse models^5,9^. However, the informative power of these DNA features is limited, and it is still challenging to anticipate *a priori* which genes are transcriptionally up- or down-regulated in RTT, raising questions whether other sequence features might also participate in transcription dysregulation mediated by the loss of MECP2^11^. Recent developments in machine learning techniques have revealed unsuspected DNA and RNA-sequence features associated with gene regulatory programs^12,13^. Employing these techniques in the RTT context could help explain the molecular mechanisms of MECP2 function and uncover other DNA sequence features important for MECP2 function.

Genome-wide analyses of steady-state mRNA levels and transcription rate changes in RTT models have demonstrated a global dysregulation of gene expression^5,9,14–18^. It is uniformly acknowledged that the magnitude of mRNA steady-state level changes is surprisingly small^3,4,14,18,19^. In 2017, Johnson *et al*^17^ used GRO-Seq for nascent RNA and RNA-Seq of chromatin, nuclear, and cytoplasmic subcellular fractions to reveal the small steady-state alterations in the *Mecp2*-null mouse brain is the result of a previously unsuspected post-transcriptional regulatory mechanism. They proposed that large transcription rate changes are compensated by reciprocally adjusting mRNA half-life, and they provided initial support for the role of two RNA-binding proteins (RBPs). In particular, they examined the enrichment of 12 RBP binding sites in subsets of mRNAs and identified HuR-(ELAVL1) *cis*-acting elements in the 3’UTR with extended mRNA stability, or AGO2 *cis*-acting elements with reduced stability to implicate the action of unknown miRNAs^17^. Their model of post-transcriptional regulation is similar to transcription buffering where RBPs present in the nucleus tag nascent mRNAs and shuttle with them to the cytoplasm. The RBPs then modify half-life to buffer transcription rate changes that preserve steady-state levels (reviewed by Hartenian E *et al*. 2019^20^). These results have not been independently tested in mouse and it is unknown whether the mechanism is conserved in human RTT neurons. Therefore, it is important to experimentally measure the direction and magnitude of half-life changes in human *MECP2*-null neurons. Moreover, the post-transcriptional mechanism has not been studied by systematic enrichment analysis of all known miRNA and RBP *cis*-acting elements of the post-transcriptionally regulated mRNAs.

Here, we simultaneously investigate the potential role of sequence features mediating transcriptional dysregulation in RTT and expand on the post-transcriptional findings of Johnson *et al* in human isogenic induced Pluripotent Stem Cell (iPSC)-derived RTT neurons. We used RNA-approach to equilibrium-sequencing (RATE-Seq) to measure transcription rate and half-life changes and employed machine learning to uncover sequence features underlying these changes in human and mouse RTT models. In parallel, we compared our human neuron findings to the high confidence RNA-seq from subcellular fractionations of *Mecp2* mutant mouse brains^9^. We found that transcription rate changes in both human and mouse datasets are best predicted by combinations of three dinucleotide frequencies in gene-bodies that include the expected CA/CG motifs, but are most accurate if they also include other dinucleotides. We discover extensive half-life changes that identify: 1) a gene set with exclusive mRNA stability dysregulation (half-life only) and no associated transcription rate changes; and 2) a larger buffered gene set in which half-life regulation compensates for transcription rate changes that fully offset or minimize mRNA steady-state changes. We demonstrate a global absolute downregulation of miRNA levels, that corresponds with a global absolute half-life increase in RTT neurons. We found individual enriched miRNA binding-sites in the half-life only gene set but very few in the buffered gene set. RBP-binding sites were enriched in the 3’UTRs of half-life only genes, and distinct sites were also enriched in buffered genes with increased transcription rate. Overall, we propose that transcription rate increases in *MECP2* neurons are subject to surveillance by RBPs that post-transcriptionally regulate RNA half-life. We find that the buffering of transcription rate changes by half-life changes is a conserved feature of RTT models which minimize the steady-state changes in mRNA levels.

## Results

### Transcription rate changes in RTT neurons do not always alter mRNA steady states

To simultaneously measure direct changes in transcription rate, half-life, and mRNA abundance levels at steady-state in RTT neurons, we performed RATE-seq on human cortical neurons derived from WT (NEU_WT_) and *MECP2*-Null (NEU_RTT_) isogenic patient-derived iPSCs^21^ (Fig 1A, Fig S1A-L). 3’end RNA-seq (QuantSeq) was utilized to quantitatively map 3’UTR isoform diversity to better understand the role of miRNAs and RBP-binding sites in half-life regulation and buffering of transcription dysregulation. Multiple RNA spike-ins controlled for 4sU pulldown efficiency, background contamination, and equivalent cell numbers. Steady-state levels were measured from an input sample taken at the 24-hour time-point. The transcription rate was measured from 0.5- and 1-hour time-points, by calculating the number of newly synthesized mRNAs in 1 hour. Finally, the half-life was derived from an equilibrium relationship between steady-state and transcription rate as a ratio of the two quantities. To measure absolute half-life in hours we estimated 4sU saturation curves as a time required to reach half of the steadystate mRNA abundance (Fig 1A, Fig S1A-L, see methods).

**Figure 1.**
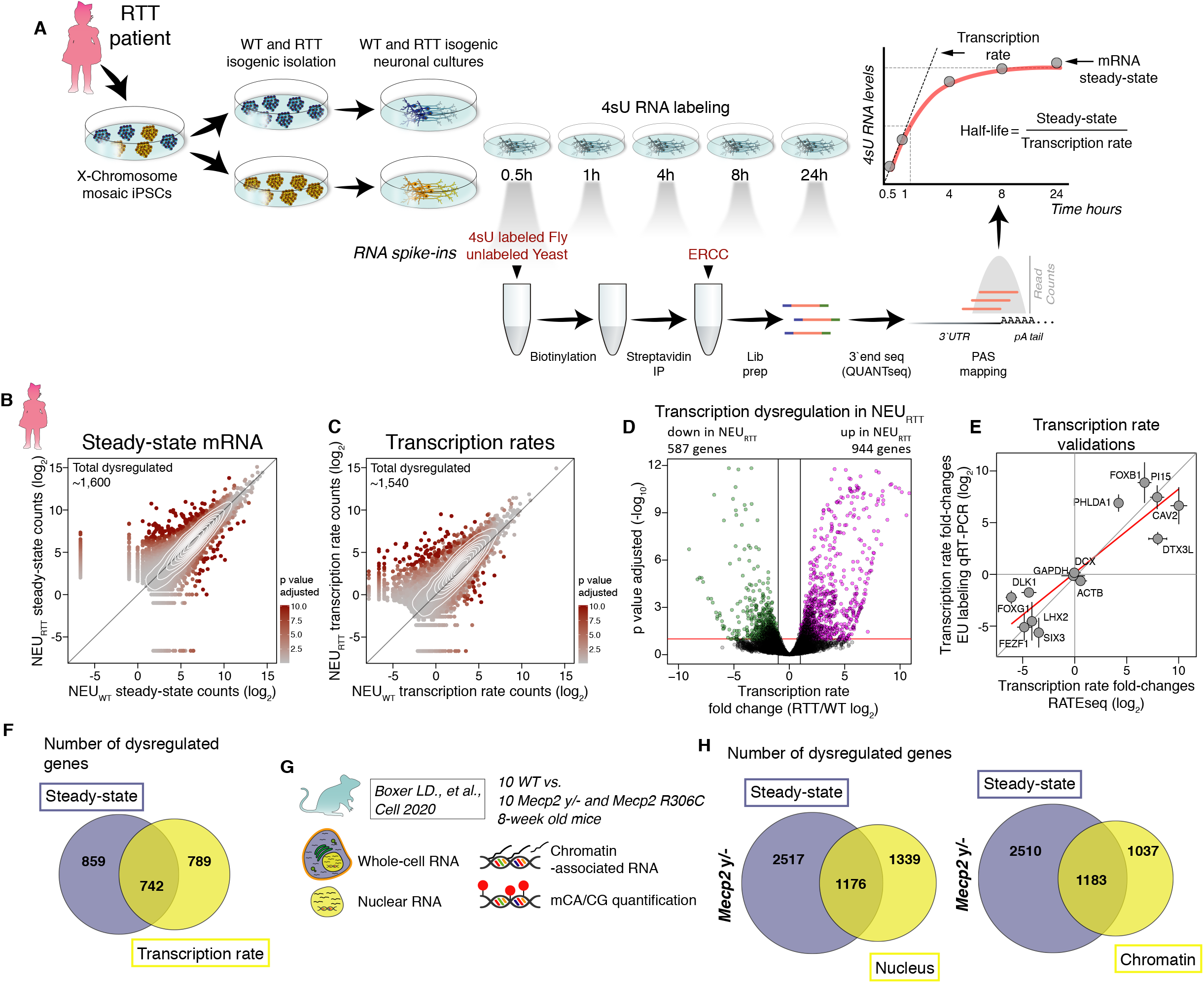
Changes in transcription rate in RTT neurons do not automatically result in altered mRNA steady states. A, schematics of experimental outline for simultaneous quantification of transcription rate, mRNA halflife, and steady-state mRNA level. An isogenic pair of human WT and *MECP2*-null iPSC-derived cortical neurons were pulse-labeled with 4sU, and at designated time-points total RNA was harvested. 4sU-labeled *Drosophila melanogaster* (fly) and unlabeled *Saccharomyces cerevisiae* (yeast), and ERCC spike-in RNAs were added as indicated and used as pull-down efficiency, non-specific binding, library preparation, and sequencing controls. Steady-state mRNA levels were quantified from an aliquot of the 24h time-point (non-biotinylated and unprocessed). B-C, scatter-plots depicting genome-wide changes in steady-state and transcription rate. D, volcano plot showing genes with increased or decreased transcription rate in *MECP2*-null neurons (NEU_RTT_). E, transcription rate fold-changes determined by RATE-seq (X-axis) were validated using an alternative approach. Neurons were incubated with 5-ethylnyl uridine (EU) and quantified following Click-it reaction and qRT-PCR (Y-axis) of genes selected to cover a large spectrum of fold-changes including genes with no changes. F, overlap of genes altered at transcription rate and/or steady-state in human NEU_RTT_. G, summary of samples from Boxer *et al* re-analyzed in our study. H, overlap of genes altered at transcription rate and steady-state in the brains of *Mecp2* y/- mouse model. Transcription rate changes in the mouse model were estimated by changes in nuclear (left Venn diagram) or chromatin-associated (right Venn diagram) mRNAs.

As expected, we found widespread dysregulation of transcription rate and steady-state in the NEU_RTT_ neurons (Fig 1B-D, S1M, Supplementary table 1). An independent 5-Ethynyl Uridine (EU) metabolic incorporation assay experimentally validated transcription rate changes of specific genes by qRT-PCR (Fig 1E). Reassuringly, comparison of our transcription rate datasets with a MECP2 ChIP-seq in mouse brain revealed that the genes with the highest changes in TR were more enriched for MECP2-binding (Fig S1N), and most transcriptionally upregulated genes displayed lower basal transcription rate in the WT controls (Fig S1O)^9^. Importantly, we found that approximately half of the transcription rate dysregulated genes in NEU_RTT_ were not altered at the steady-state mRNA level (Fig 1F).

Given the relevance of these findings to disease mechanisms, we validated the discrepancy between transcription rate and steady-state changes in an orthogonal *in vivo* RTT system. We re-analyzed the high-confidence datasets from Boxer *et al*^9^ that sequenced nuclear and chromatin-associated mRNA abundance as a proxy for transcriptional dysregulation and the whole-cell fractions from cortical forebrain samples of WT, *Mecp2 y/-*, and point-mutant *Mecp2 R306C* adult mice (Fig 1G). Our analysis shows a similar pattern where approximately half of the genes transcriptionally dysregulated are not altered at the steady-state level (Fig 1G-H). This confirms that transcription rate dysregulation in the absence of MECP2 does not automatically result in altered steady-state mRNA levels, and that unaltered steady-state mRNA level does not automatically mean there is no change in transcription rate.

### The direction of transcription rate changes in RTT neurons is predicted by gene-body dinucleotide frequencies

The high fraction or number of mCA/mCG in longer genes have been associated with the small magnitude upregulation of these genes in *Mecp2* mouse models^3,7,9^. However, consistent with Johnson *et al*^17^, our analyses did not show a gene-body length effect in genes dysregulated at the transcription rate or steady-state (Figure S2A). The contribution of other sequence features that predict which genes will be transcriptionally dysregulated in RTT models has not been systematically evaluated *in vivo*^11^. To learn sequence features of the differentially expressed genes that are relevant for the direction of transcriptional shifts in our immature RTT neurons, we trained a classifier model using our measurements of genomewide transcription rate changes in NEU_RTT_ (Figure 2A and S2B). As anticipated by the lack of correlation between gene-body length and transcription rate changes (Figure S2A), this model also resulted in random predictions based on gene-body length (including introns) (Figures 2B and S2C-D). In contrast, frequencies of dinucleotides in the gene-body produced high prediction accuracies similarly found when using either the coding sequence (CDS) or 3’UTR (Figures S2C-D). The prediction accuracies of CA/CG in gene-bodies were lower than predictions based on any combination of dinucleotides even when combined with gene-body length. In fact, removal of CA/CG from the model had no negative effect on prediction accuracy. This supports a prominent role for additional gene-body dinucleotide combinations in modulating transcription rates (Figures 2B and S2C-D).

**Figure 2.**
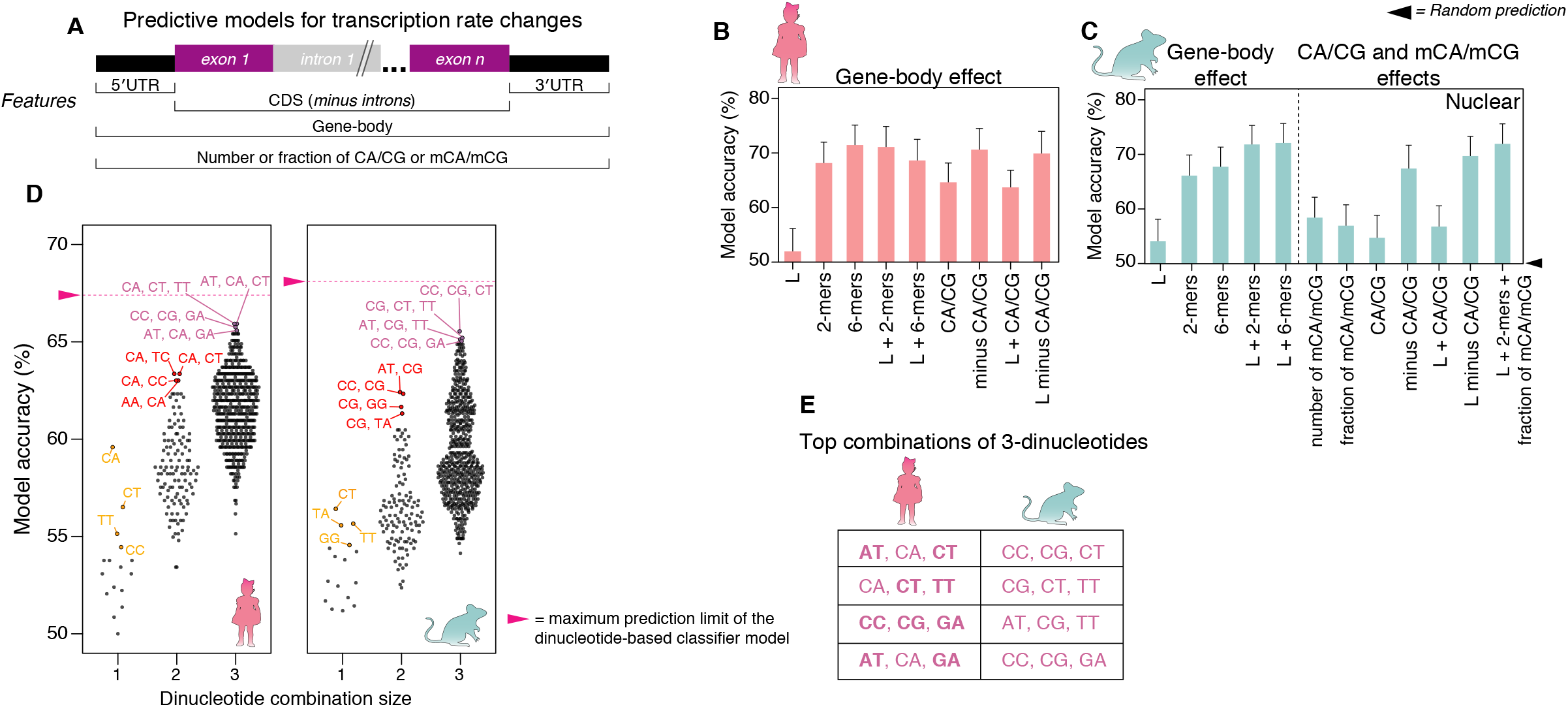
The direction of transcription rate changes in RTT neurons is predicted by three combined gene-body dinucleotide frequencies. A, random forest classifier for prediction of gene-body sequence features relevant for transcription rate fold-changes in human NEU_RTT_ and in cortical brain samples of the mouse *Mecp2 y/-*. B, percent accuracy (Y-axis) of transcription rate fold-change predictions in human NEU_RTT_ (B) or mouse *Mecp2 y/-* (C) considering different gene-body sequence features. L= gene-body length; 2-mers and 6-mers= 2 and 6-nucleotide sequence elements, respectively; CA/CG= number of CA or CG di-nucleotides; minus CA/CG= removal of CA and CG di-nucleotides from predictive models; mCA/mCG= methylated CA and CGs; + sign denotes combinations of two or more sequence features. D, top combinations of dinucleotides contributing to the full prediction accuracy described in panels B and C. Accuracy increases to the maximum when a specific combination of three dinucleotides is used. Similar behavior is observed between human (left) and mouse (right) datasets. E, top combinations of three dinucleotides contributing to the full prediction model. Predictive di-nucleotides conserved between human and mouse are denoted in bold fonts in the human column.

To test the combined dinucleotide-based predictive model in an orthogonal *in vivo* system, we trained a classifier model on the data from Boxer *et al* which also includes cytosine methylation quantification (Fig 1G)^9^. In the adult mouse brain, our model captured gene-body length and mCA fraction as predicting whether a gene is transcriptionally up-regulated in the absence of *Mecp2* (Figures S2E-F). However, these models do not discern whether a gene is transcriptionally down-regulated, nor discriminate up-versus down-regulated genes (Figures S2E-F). In contrast, the gene-body dinucleotide frequencies captured the direction of most transcriptional dysregulation in the *Mecp2* y/- mouse, with lower accuracy predicted by the CDS and UTR separately, an effect that was independent of gene-body length (Figures 2C and S2G-I). Surprisingly, combining CA/CG frequencies had less predictive accuracy than combinations of remaining dinucleotides independent of their methylation status (Figures 2C, S2G-I). To investigate which dinucleotide or combinations thereof were responsible for the high predictive accuracy of transcription rate changes in RTT neurons, we repeated the classifier model considering single or multiple dinucleotide combinations. The classifier found that specific combinations of three dinucleotides reached the predictive accuracy of the full model for both human and mouse (Figure 2D). The top four combinations of three dinucleotides were highly similar between human and mouse, and included AT as enriched along with CA and CG (Figures 2D-E) and other dinucleotides. Taken together, the classifier models of fetal stage human neurons and adult mouse brain indicate that combined frequencies of three dinucleotides that include non-CA/non-CG dinucleotides contribute to the direction of transcription rate modulation mediated by MECP2.

### mRNA half-life changes in RTT neurons directly alter the steady-state or buffer transcription rate

The RATE-Seq half-life measurements revealed widespread changes in mRNA stabilities that caused a global absolute increase in the mean half-life in NEU_RTT_ (Figures 3A-B, S1A-L, and Supplementary table 2). These measurements were independently validated on specific genes using transcription inhibition and qRT-PCR (Figure 3C). We found approximately 860 genes (~45% of all genes with altered half-life) with significantly increased or decreased mRNA half-life but with unchanged transcription rate, termed half-life only (HL-only), leading directly to changes in the steady-state levels (Figures 3D and S3A. See also Figure 1E). These analyses show that a significant fraction of genes dysregulated at the steady-state level were exclusively driven by changes in mRNA stability.

**Figure 3.**
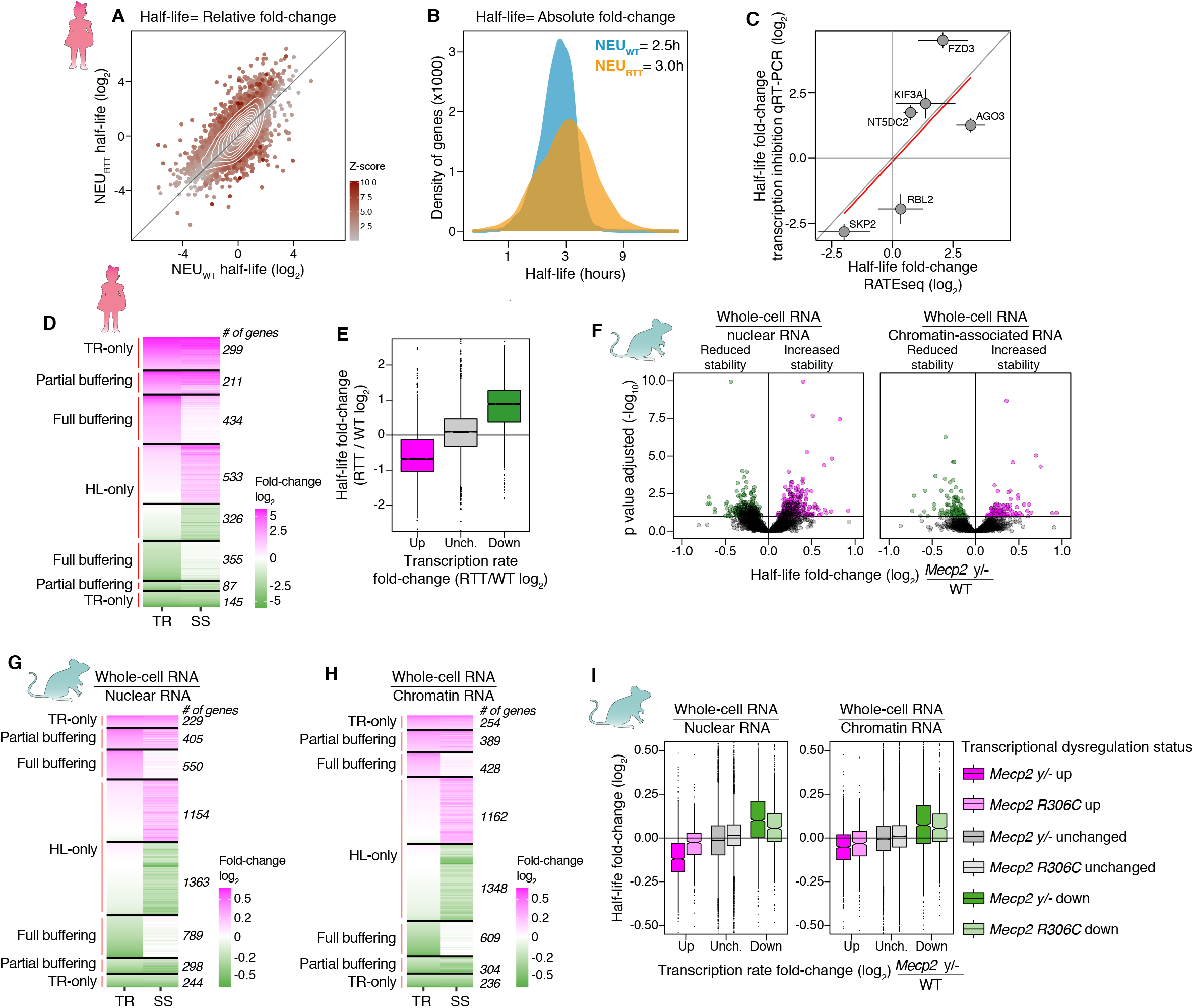
Widespread changes in mRNA half-life directly alter steady-state abundance or buffer transcription rate changes in human and mouse RTT models. A, scatter-plot showing global relative changes of half-life in NEU_RTT_. B, global increase in the absolute mean half-life in NEU_RTT_. C, half-life fold-change determined by RATE-seq (X-axis) were validated using an alternative approach. Neurons were incubated with Actinomycin D (transcriptional inhibitor) and RNA harvested at different time-points. Selected genes covering a wide range of fold-changes were quantified by qRT-PCR (Y-axis). D, number of genes with changes in transcription rate only, partially or fully buffered by half-life changes, and genes whose change in steady-state are caused by altered half-life only. Genes were defined with transcription rate only changes when the difference between steady-state and transcription rate log2 fold-change was less than 25% of transcription rate log2 fold-change. E, most genes with increased (magenta box) or decreased (green box) transcription rate show decreased or increased half-life, respectively, and genes with unchanged transcription rate do not display significant changes in half-life as a group. F, global changes in half-life detected in brains of the *Mecp2* y/- mouse model estimated from both nuclear (left) or chromatin-associated RNAs (right) (horizontal line denotes FDR = 0.1). G-H, number of genes with transcription rate changes that are partially or fully buffered by mRNA stability mechanisms, and genes whose change in steady-state are caused by altered half-life only in the mouse model. I, the *Mecp2* y/- mouse models also display the reciprocal behaviour in half-life regulation relative to transcription rate. TR= transcription rate, HL= half-life, SS= steady-state

We then explored the impact on steady-state mRNA levels when both half-life and transcription rate are changed. We found that in these cases, half-life moved in the opposite direction to transcription rate and decreased the net change in steady-state levels (Figures 3D-E). We found that 789 genes exhibited full buffering where the RNA half-life entirely offset the transcription rate change resulting in no steadystate change. A further 298 genes showed half-life changes that partially counteracted the transcription rate changes modulating steady-state changes. The contribution of half-life to the steady-state level changes (Figure S3B) and buffering (Figure S3C) was independent of the fold-change or FDR thresholds chosen. None of the measured changes correlate with gene-body or processed transcript lengths (Figures S2A and S3D). We found ~28% of all genes (n= 444) with altered steady-state expression to be exclusively dysregulated at the transcription rate level (TR-only), underscoring the role of post-transcriptional regulation of mRNA stability in directly altering or buffering the RTT transcriptome.

#### Half-life shifts and transcription buffering are conserved in *Mecp2* mouse models

Given the importance of finding novel global changes in half-life in the NEU_RTT_ neurons and its implications for interpreting steady-state level dysregulation typically found in RTT models, we investigated whether these results were reproducible *in vivo*. To determine half-life changes in the mouse brain in RTT models we re-analyzed the data from Boxer *et al* for whole-cell, nuclear, and chromatin-associated RNAs^9^ to determine changes in half-life. We accomplished this by comparing the abundance of transcribed genes in the nucleus and chromatin fractions, to the whole-cell fraction that includes mRNA undergoing decay in the cytoplasm^17^ (see methods). From the 5,032 genes reported by Boxer *et al* to be differentially regulated at steady-state in the *Mecp2* y/- mice, we found a similar pattern of widespread changes in half-life (Figure 3F, Supplementary table 3). We also found similar proportions of HL-only genes and the extent of full or partial buffering in the mouse models (Figures 3G-I and S3E. See also Figure 1H). In line with our RTT human neuron findings, the contribution of half-life to the steady-state level changes (Figure S3F) and buffering (Figure S3G) was also independent of the fold-change or FDR thresholds chosen.

Despite the conserved relationship that half-life and transcription rate have on steady-state gene expression in human and mouse RTT models, we found minimal overlap in the identities of genes dysregulated in each species (Figures S3H-I). Furthermore, minimal overlap was also observed between the *Mecp2 y/-* and *Mecp2 R306C* mouse models as already observed by Boxer *et al* (Figures S3J-L). Overall, our results consistently identify steady-state level changes driven by half-life only without any measurable transcription rate shift in both human and mouse RTT models. Importantly, more than half of all half-life shifts fully or partially buffer *Mecp2*-mediated transcription rate dysregulation, and only a small number of genes have increased steady-state changes due to combined half-life and transcription rate changes in the same direction. Finally, similar to the human findings, we only find 473 genes (13% of total genes altered at steady-state, which excludes the full buffered group) with transcription rate only dysregulation as measured in the nuclear fraction. Altogether, our findings demonstrate a pattern of halflife shifts and transcription buffering that is conserved in RTT mouse and human models.

### Cis-acting elements in the 3 UTRs are highly associated with half-life changes

Our findings indicate that RNA half-life is a critical regulatory layer defining steady-state RNA levels in RTT models. We therefore considered several potential mechanisms underlying how RNA halflife is controlled in RTT neurons: 1) alternative polyadenylation; 2) alternative-splicing; 3) codon usage integrating translation elongation to RNA stability; 4) sequence composition of 3’UTR and gene-bodies; and 5) enrichment of miRNA binding-sites and RBP *cis*-acting elements in the 3’UTR between buffered and non-buffered genes. Initially we investigated the contribution of mRNA isoforms by mapping and quantification of 3’UTR alternative poly-Adenylation events that showed no difference in the frequency of poly-Adenylation site usage in NEU_RTT_ (Figures S4A-C). Moreover, measurement of 3’UTR (NEU_WT_ vs. NEU_RTT_) and alternatively-spliced mRNA isoforms (mouse Boxer *et al*) indicated that all mRNA isoforms display the half-life buffering effect to the same degree (Figures S4D-F). These analyses argue against changes in mRNA isoform usage participating in the half-life shifts in RTT models.

Next, we created a new classifier predictive model to estimate the effect of codons and sequence composition on the direction of changes to mRNA half-life (Figure 4A). Overall 3’UTR length and nucleotide frequency had no predictive value for classifying increased or decreased half-life changes (Figure 4B). Transcription rate changes were anti-correlated with half-life leading to high predictive accuracy. These results underscore the unidirectional and reciprocal link between transcription rate dysregulation and compensatory mRNA stability control in NEU_RTT_. Dinucleotide frequencies in the 3 ‘UTR offered significant prediction accuracy on whether the half-life was increased or decreased. Increasing the size of the tested k-mers from dinucleotides to 4-mers and 6-mers to encompass potential *cis*-acting elements improved the prediction accuracy of half-life (Figures 4B and S4G), equivalent to transcription rate alone. A combination of transcription rate and 6-mers showed no further improvement in accuracy. Curiously, we found that classifier features in the CDS mirror that of the 3’UTR models also offering significant prediction accuracies for the half-life changes (Figures S4H-I), highlighting a significant sequence composition correlation between CDSs and 3’UTRs (Fig S4J). In contrast, comparison of the prediction models for in-frame codons (3-nt sequences) indicates that codon optimality has no effect on half-life changes in NEU_RTT_ (Fig S4I). Importantly, while the predictive accuracy of k-mers is lower in mouse, the classifier model predictions are upheld in the *Mecp2* mouse model. These findings show a reproducible effect of transcription rate on half-life, thereby implicating a conserved buffering mechanism through *cis*-acting elements impacting half-life (Fig 4C, S4 L-N).

**Figure 4.**
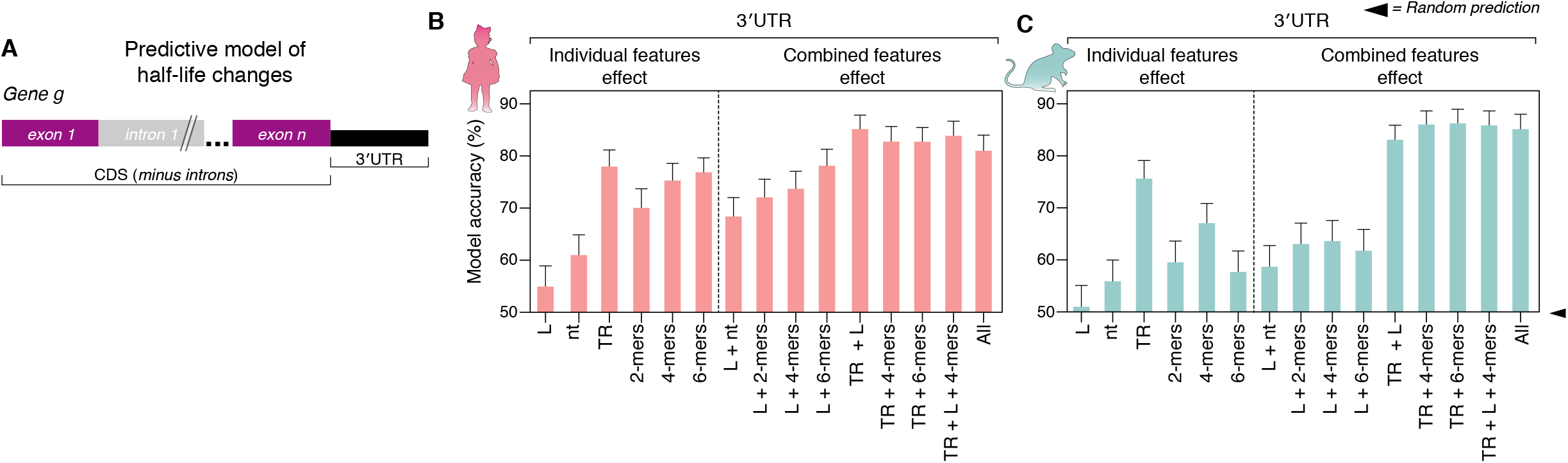
*Cis*-acting elements in the 3’UTR are highly associated with half-life changes. A, random forest classifier for prediction of mRNA sequence features relevant for half-life fold-change in human NEU_RTT_ and in cortical brain samples of the mouse *Mecp2 y/-*. B, percent accuracy (Y-axis) of half-life fold-change predictions in human NEU_RTT_ (B) or mouse *Mecp2 y/-* (C) considering different mRNA sequence features. >80% prediction accuracies can be achieved with the features tested, and indicate that 3’UTRs contain sequence elements relevant for half-life changes in both humans and mice. L= gene-body length; nt= nucleotide sequence; 2-mers, 4-mers, and 6-mers= 2, 4, and 6-nucleotide sequence elements, respectively; TR= transcription rate; All= all features considered at the same time; + sign denotes combinations of two or more sequence features.

### miRNA and RBP cis-acting elements correlate with half-life changes in RNA stability exclusive genes

To examine a possible role of miRNAs in mRNA half-life regulation, we first performed small RNA-seq with a spike-in strategy to inform on relative and absolute changes in miRNA abundance between the isogenic human NEU_WT_ and NEU_RTT_. These results showed that the steady-state levels of miRNAs changed by as much as 4-fold up or down (Figure 5A, Supplementary table 4). Interestingly, most changes in miRNA steady-state abundance were captured in the RATE-Seq data by shifts in the transcription rate of miRNA genes (Fig 5B, Supplementary table 4). These findings indicate that transcription dysregulation of miRNA genes drives the changes in miRNA abundance, in addition to miRNA maturation processing as indicated previously^22,23^. Moreover, after normalization against the spike-in RNAs, the scaled absolute fold-change of most miRNAs was decreased in NEU_RTT_ (Fig 5C-D). This global decrease in miRNA abundance in NEU_RTT_ is in line with the global absolute upshift of median half-life (Figure 3B).

**Figure 5.**
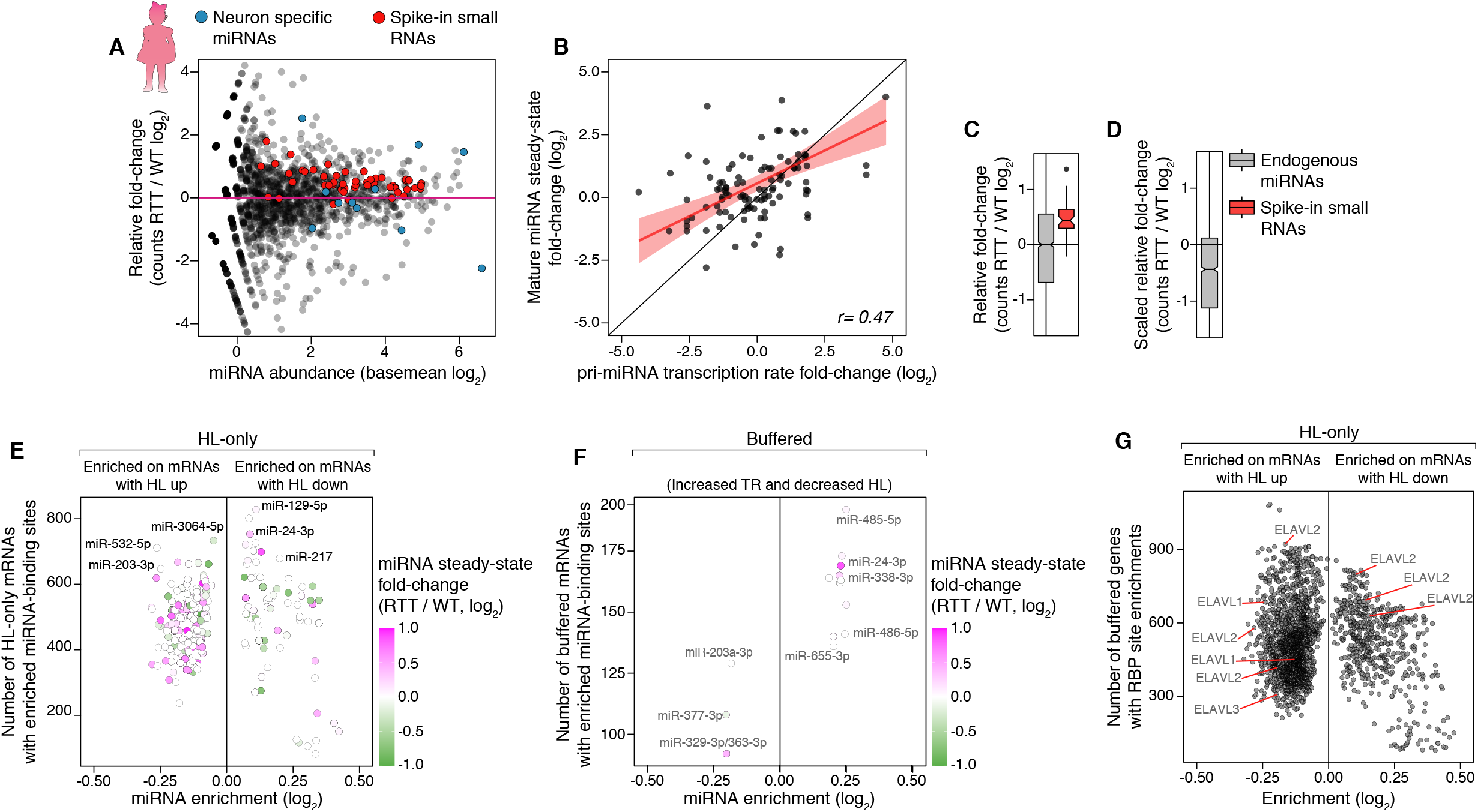
miRNA and RBP *cis*-acting elements correlate with half-life changes in RNA stability exclusive genes. A, small RNA-seq to quantify changes in human miRNA abundances in the NEU_RTT_. X-axis represents the basal abundance of each mature miRNA detected in the NEU_WT_ and Y-axis represents their foldchange in the NEU_RTT_. Blue dots, neuronal-specific miRNAs showing that they accumulate at abundances higher than the mean in both NEU_WT_ and NEU_RTT_. Red dots, the abundance of the small RNA spike-ins added on a per-cell basis and used for library preparation control and quantification of miRNA abundance at the absolute levels. B, comparison of the fold-changes between steady-state mature miRNA levels and primary miRNA (pri-miRNA) transcription rate in NEU_WT_ and NEU_RTT_ showing a significant correlation between both (*r*= *0.47, p val 4.3^-7^*), indicating many changes in mature miRNA steady-state levels are caused by changes in their transcription rate. C, DESeq2 miRNA steady-state level fold-change before spike-in normalization, and D after normalization based on the spike-in fold-change. The absolute abundance of miRNAs in NEU_RTT_ is reduced for most miRNAs. E-F, miRNA-binding sites enriched in half-life only (panel E) or in combination with transcription rate up and half-life down (buffered, F). Y-axis represents the number of genes containing each of the miRNA-binding sites found, and X-axis the enrichment levels. Color represents the miRNA steady-state fold-change of the respective miRNA in the NEU_RTT_ as measured in A. G, 7-mers known to be targeted by RBPs enriched in the HL-only group based on the RNAcompete database^25^. TR= transcription rate, HL= half-life.

To investigate the role of these miRNA changes in the regulation of half-life in NEU_RTT_, we performed motif enrichment analysis of miRNA-binding sites present in the TargetScan database^24^. This analysis identified multiple potential miRNAs-binding sites as enriched in up to 800 genes in the HL-only group of mRNAs whose steady-state level changes are exclusively directed by half-life changes (Figure 5E, Supplementary table 5). In contrast, we found significantly fewer miRNA-binding sites sequences enriched in the group of buffered mRNAs and many of these show no change in the miRNA abundance (Figure 5F, Supplementary table 5). Overall, these data indicate the reduced miRNA abundance contributes to the regulation of HL-only genes in NEU_RTT_. However, very few individual miRNAs correlate with buffering, although combinatorial effects of multiple miRNAs cannot be excluded.

We then performed an unbiased search for enrichment of 174 RBP *cis*-acting elements, as described in the RNACompete database^25^, in the 3’UTR of mRNAs with altered half-life in NEU_RTT_. We found hundreds of enriched RBP targets in the group of HL-only genes (Figure 5G, Supplementary table 5). This result includes the RBP HuR (ELAVL1) whose different motifs are enriched in 300-900 mRNAs with increased half-life in NEU_RTT_. Our results suggest that 3’UTR-directed miRNA and RBP regulation best explain the HL-only gene set changes.

#### Only RBP *cis-acting* elements are enriched in buffered genes with increased transcription rate and decreased half-life

Having excluded miRNAs as playing a substantial role in mRNA buffering, we explored a role for the 174 RBP *cis*-acting elements in the buffered group of mRNAs. We observed that no RBP *cis-acting* elements were enriched for genes with decreased transcription rate and increased half-life (Figure 6A, right panel), and only a handful were depleted. This result suggests that RBPs are not involved in regulating transcripts with decreased transcription rate for half-life stabilization. In contrast, we found numerous RBP elements that were enriched in 100-200 buffered genes with increased transcription rate and decreased half-life (Figure 6A, left panel and Supplementary table 5). To identify RBPs that plausibly regulate nascent mRNAs, we aggregated all the buffering enriched RBPs with reported cellular localizations, and found that 51 were nuclear and/or able to shuttle to the cytoplasm whereas only 18 were reported to be cytoplasmic only (Figure 6B). To examine roles of specific RBPs, we first noted that ELAVL1 elements are depleted in >150 genes in this set, demonstrating that it is only enriched in the HL-only gene set. In contrast, *NOVA2* and *ZFP36* were enriched in more than 200 buffered genes and these RBPs bind premature mRNA co-transcriptionally^26,27^. Additionally, *PTBP1* and multiple arginine-serine rich (SRSF) splicing factors that have been shown to participate in transcriptional buffering^28^ were enriched in >150 genes. Further support for a directional role by *NOVA2* and SRSFs is that they were depleted in the buffered genes with decreased transcription rate and increased half-life (Figure 6A). Our results reveal specific sets of RBP motifs associated with half-life regulation of buffered genes with increased transcription rate in RTT neurons.

**Figure 6.**
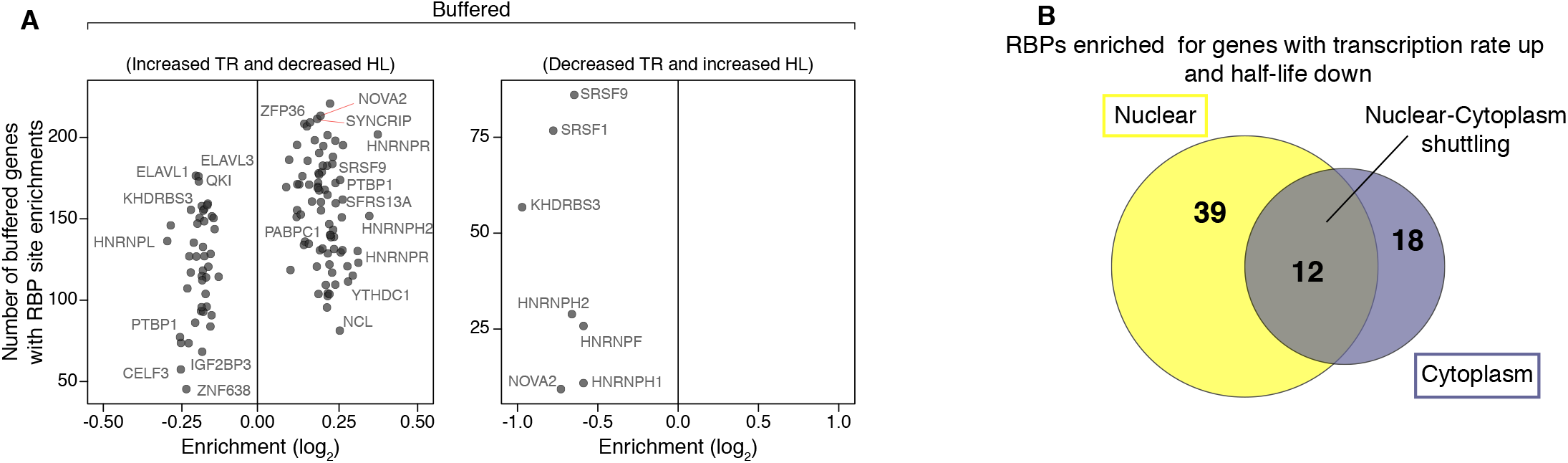
RBP *cis*-acting elements are enriched in buffered genes with increased TR and decreased half-life. A, 7-mers known to be targeted by RBPs enriched in the group of buffered mRNAs. A different set of RBP *cis*-acting elements was found enriched compared to the HL-only group (Figure 5G). B, cellular distribution of the RBPs enriched in mRNAs with increased transcription rate and decreased half-life showing that these are predominantly nuclear.

## Discussion

Our analysis of a new dataset of transcription rate and half-life changes in human neurons and independent re-analyses of mouse RTT models demonstrate that most transcription rate changes in the absence of *MECP2* are buffered by post-transcriptional regulation of mRNA stability. We used RATE-Seq to simultaneously measure transcription rate and half-life changes in human neuron samples. We complemented this approach by applying the subcellular fractionation strategy employed by Johnson *et al* to the mouse resource dataset of Boxer *et al*. The consistent findings from both methods and model systems unambiguously show that post-transcriptional regulation is a modifier of transcription rate dysregulation in RTT, and that it is a conserved mechanism shared by human and mouse neurons. We provide evidence for changes in steady-state levels driven solely by half-life only shifts, and large transcription rate shifts that are entirely offset at the steady-state level by half-life mechanisms that we refer to as buffering. These observations have major implications for interpreting RNA-Seq results in RTT and potentially other neurodevelopmental disorders, or in diseases of other tissues caused by mutations in genes that modulate transcription like *MECP2*. The existence of transcriptional buffering mechanisms in mammals raises a cautionary note for interpreting RNA-Seq steady-state results in the general context of transcriptional regulation. It also argues in favour of more widely prioritizing methods that directly measure nascent transcription or account for mRNA stability. Moreover, we extend the limited search by Johnson *et al* who found two RBP *cis*-acting elements enriched in mRNAs with altered half-life by discovering hundreds of new RBP *cis*-acting elements in the half-life only gene set. We also demonstrate the enrichment of miRNA-binding sites in the HL-only gene set. The buffered gene set with increased transcription rate likely describes the genes transcriptionally repressed by MECP2^29^. In this group, we identified a restricted subset of mostly nuclear or shuttling-capable RBPs whose *cis*-acting elements are highly enriched. Our findings thus reveal numerous candidate RBPs potentially involved in buffering the increased transcription rate of their mRNA targets. This mechanism may act as a network to coordinate mRNA degradation in healthy neurons and to compensate for transcription rate dysregulation in RTT neurons.

Our computational methods were focused on defining *cis*-acting elements that are relevant for transcription rate or half-life regulation. At the DNA level, our classifier model discovered that the direction of transcription rate shifts in human and mouse RTT models are best predicted by combinations of three dinucleotides that include the canonical MECP2-binding sites CA and CG, together with other dinucleotides including AT. The MECP2 AT-hook domain contributes to low-affinity transient interactions with AT-rich DNA that influence the dynamics of MECP2 binding to local high affinity methylated DNA sites^30^. Tellingly, the classifier predictions are low when modelling only CA/CG dinucleotide frequencies without accounting for low-affinity sites, and the accurate predictions are unaffected when low-affinity sites are included but the CA/CG dinucleotide frequencies are omitted. These unbiased findings from machine learning algorithms indicate that the gene-body frequencies of other dinucleotides like AT are important and are a conserved mechanism defining the direction of transcription rate changes in RTT neuronal models. We speculate they may act by transient recruitment of MECP2 influencing its local binding dynamics to nearby methylated DNA. We confirmed a role of gene-body length in transcription rate regulation but only in the adult mouse neuron resource of Boxer *et al*. However, neither our classifier model on the human neuron RATE-Seq dataset nor the adult mouse dataset from Johnson *et al* support this observation. It is possible that gene-body length contributes less to transcription rate regulation in fetal stage neurons derived from iPSC in which only mCG modifications are expected to be present^7,31^, or that it requires the power of ten replicate samples used in the Boxer *et al* resource to be detected.

With regards to half-life only regulation, to define which miRNA-binding sites to investigate we first used our human neuron miRNA dataset to identify the miRNAs that are altered in NEU_RTT_. These results confirm the reported miRNA changes in RTT mouse models, although the RATE-Seq dataset shows that intergenic pri-miRNA transcription rate is altered in RTT adding another dimension to the known miRNA processing alterations in mouse^22^ and human neurons^23^. Through the use of spike-in scaling in the miRNA dataset, we deduced a global absolute downregulation of miRNAs in NEU_RTT_ that account for the global absolute increase in mRNA half-life of ~0.5 hrs. Superimposed on the increased global half-life effect were individual HL-only genes which our classifier models revealed strong enrichment of miRNA-binding sites consistent with their known role in mRNA instability. Many RBP *cis*-acting elements including ELAVL1 (HuR) were enriched in the HL-only gene set. While Johnson *et al* reported HuR and AGO2 *cis*-acting element enrichments in buffered gene sets in the mouse using the subcellular fractionation approach, our RATE-Seq results and unbiased search of RBP *cis*-acting elements point to a role for miRNAs and ELAVL1 in mRNA stability in human neurons rather than in the buffering mechanism itself.

While we eliminated several possible buffering mechanisms, one limitation is that we were unable to test a potential role of mRNA methylation modifications or poly(A)-tail length regulation on the targeted mRNAs with altered transcription rate^32^. We speculate that these mechanisms could also participate in the buffering mechanisms, particularly in the gene set with decreased transcription rate and increased half-life that we found was not associated with enrichment of either miRNA-binding sites or RBP *cis*-acting elements. The simplest mechanism for buffered genes with increased transcription rate is that the RBPs bind nascent mRNA in the nucleus and are transported to the cytoplasm where they tag the transcript for stabilization. We speculate that the nuclear RBPs are limiting in neurons, and if transcription rate increases then the proportion of tagged mRNA falls, leading to relatively more degradation in the cytoplasm and decreased half-life. A more complex variation on this model is that the concentration of some RBPs themselves may also change in RTT, and this may increase or decrease their ability to stabilize their target mRNAs. To distinguish these models, it will be necessary to determine which RBPs are changed at the protein level in RTT using proteomics of neuronal nuclei and then individually testing their impact through gain- or loss-of-function assays on the mRNA targets.

Equivalent loss-of-function experiments have already been described in humans with neurodevelopmental disorders caused by mutations in RBP genes such as *NOVA2*^27^. The impact of these RBPs on buffering could be established using existing or new iPSC or mouse models. In fact, global transcription rate and RBP concentrations will inevitably be altered during the course of neurodevelopment, suggesting that it would be valuable to define the buffered gene sets in iPSC and their progeny Neural Progenitor Cells relative to the final neurons described here. We and others^18,33^ have previously reported translational regulation changes in RTT neurons in both ribosomal loading and protein stability implemented through alterations of E3-ubiquitin ligase protein levels. These findings emphasize that buffering in RTT and potentially other disorders likely operates at both the mRNA and protein levels.

## Acknowledgments

This study was funded by grants from the Canadian Institutes of Health Research (CIHR; PJT-148746, PJT-168905 to J.E.); the Canada First Research Excellence Fund (Medicine by Design Cycle I: J.E.); the Col. Harland Sanders Rett Syndrome Research Fund at the University of Toronto (J.E.); the Ontario Brain Institute (POND Network: J.E.); and John Evans Leaders Fund/Canada Foundation for Innovation (JELF/CFI: J.E); Canada Research Chairs Program (M.D.W. and J.E.); Early Researcher Award from the Ontario Ministry of Research and Innovation (M.D.W.); Genome Canada Disruptive Innovation in Genomics Grant (M.D.W to support K.E.Y.); NIH grant R35GM128680 and the University of Colorado RNA Bioscience Initiative (O.R.); David Steven Cant Scholarship (M.M.) We thank the Centre for Applied Genomics (TCAG) at SickKids for RNA sequencing, and Brian Kalish for comments on the manuscript.

## Author contributions

Conceptualization, D.C.R., M.M., A.N., O.R., and J.E.; Investigation, D.C.R., M.M., A.N., K.Y., W.W., A.P., and J.L.; Software, M.M., and A.N.; Writing – original draft, D.C.R., M.M., and J.E.; Writing - Review & Editing, D.C.R., M.M., A.N., K.Y., P.P., O.R., M.W., and J.E.; Visualization, D.C.R., and M.M.; Supervision & Funding Acquisition, O.R., M.W., and J.E.

---

**Fig S1.**
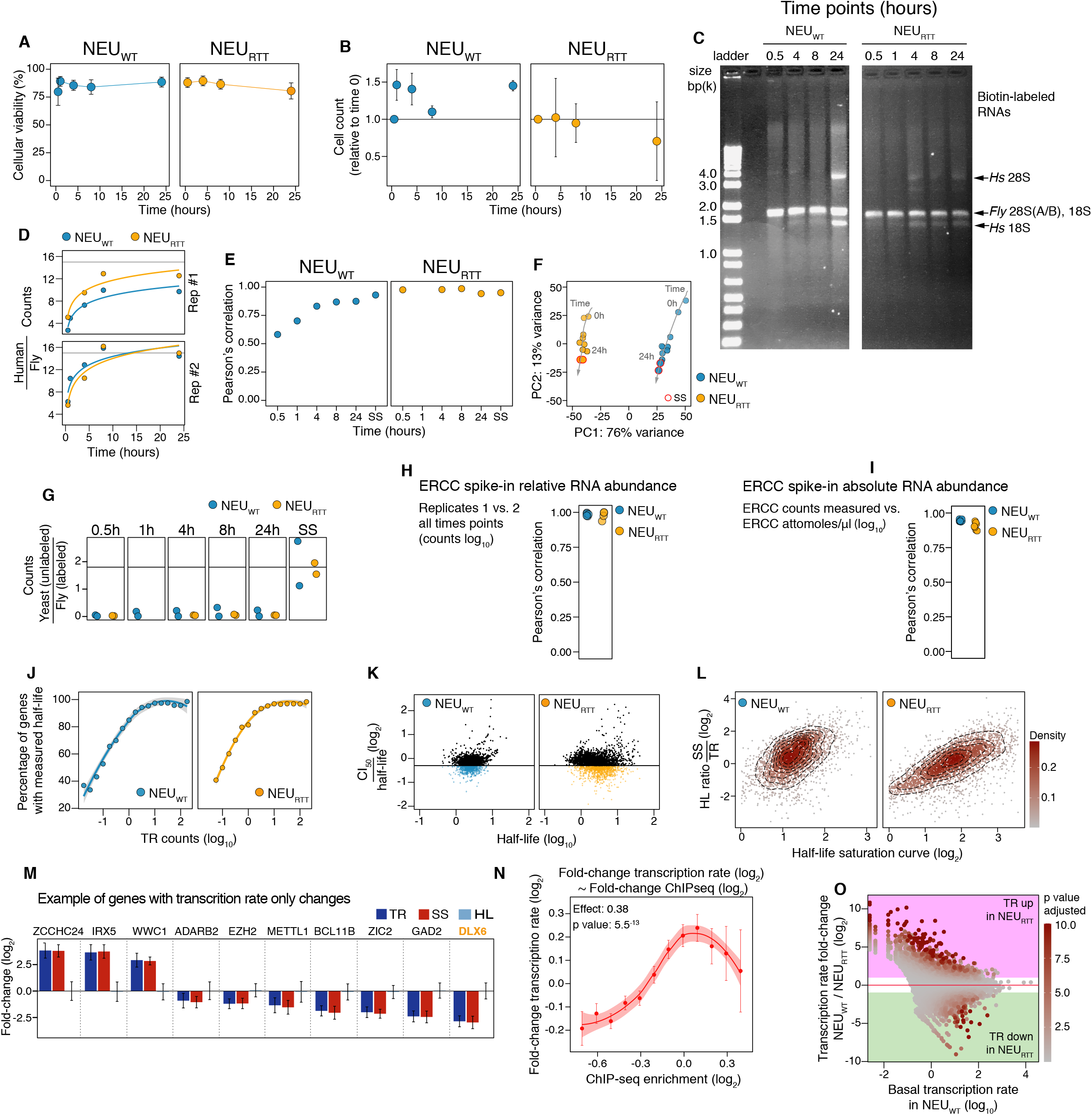
Quality control of the RATE-seq experiment. A-B, 4sU does not cause changes in neuronal viability or cell numbers at the dose and times used in the experiment. C, representative agarose gel showing 4sU-labeled pulled-down RNA for each time-point. The presence of human (*Hs*) and fly ribosomal RNAs are denoted by arrows. D, 4sU incorporation kinetic curves of human mRNA normalized to fly spike-in RNA. E, Pearson’s correlation between replicates of the human RNA for each time-point. F, principal component analysis of sequencing data from NEU_WT_ and NEU_RTT_ showing significant separation between genotypes and time-points. Steady-state samples are denoted with the red outline, and cluster the closet with the 24-hour time point samples from where they are derived. G, read counts of spike-in RNAs: unlabeled yeast relative to fly RNA for all time points in NEU_WT_ and NEU_RTT_. Unlabeled yeast RNA spike-in RNA was used as pull-down negative control (background control). The absence of yeast RNA indicates that the streptavidin-biotin pull-down of 4sU labeled RNAs had minimal contamination of unlabeled human RNAs, but were readily detectable in the steady-state samples. H-I, Pearson’s correlation of ERCC spike-in RNAs between (H) replicates, and (I) spike-in concentration and sequencing measured abundance used for all time points in NEU_WT_ and NEU_RTT_. ERCC spike-in RNA was used as control for library prep quality. High Pearson’s correlations indicate high-quality of the library samples. J-K, half-life measured with 4sU saturation curve method. (J) percentage of genes with measured half-life depending on transcription rate. K, accuracy of half-life shown against half-life magnitude. CI_50_ stands for 50% confidence interval. Black points denote genes with poorly fit saturation curves and are removed from analysis in panel L. L, comparison of half-life estimated with two methods for well measured genes selected in panel K. M, example of genes displaying transcription-mediated changes in steady-state (TR-only). DLX6 is known to confer high risk for autism when mutated. N, human genes with the highest transcription rate fold-change are also enriched for MECP2 binding as previously described in mice^9^ by MECP2 ChIP-seq. O, genes with the highest increase in transcription rate fold-change in NEU_RTT_ had lowest basal transcription rate in NEU_WT_ neurons as seen previously^17^. TR= transcription rate, HL= half-life, SS= steady-state.

**Figure S2.**
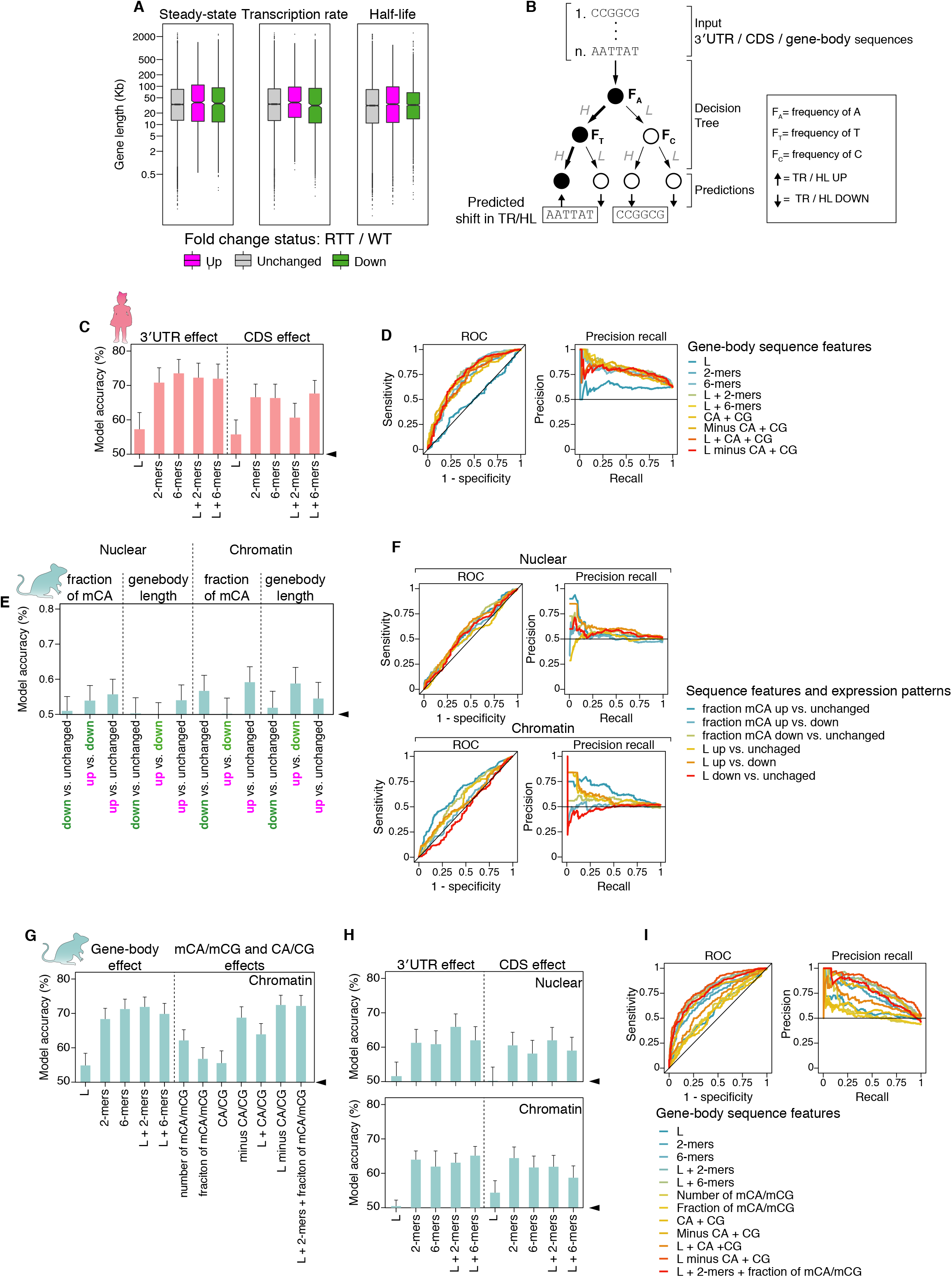
Classifier model to predict DNA sequence features affecting transcription rate in the absence of *MECP2*. A, human genes with steady-state, transcription rate, or half-life fold-changes in NEU_RTT_ do not show significant enrichments for gene length as seen for adult mice^9^. B, schematic representation of a decision tree for the classifier utilized. A collection of decision trees forms the random forest model. C, 3’UTR or CDS mRNA sequence features-based random forest classifier for prediction of transcription rate foldchange in NEU_RTT_. D, receiver operating characteristic curve (ROC) and precision-recall curve (PRC) showing overall performance of the classifier for prediction of human transcription rate fold-changes. E, predictive model for transcription rate fold-changes in the mice using either nuclear or chromatin-associated RNA samples. While the fraction of mCA and gene-body length offer some accuracy to distinguish transcription rate up versus unchanged or up versus down in some cases, the classifier found that other sequence features offer higher prediction accuracies (see also figure 2C). F, ROC and PRC showing overall performance of the classifier for prediction of mouse transcription rate fold-change relative to panel E. G, gene-body or number and frequencies of mCA/mCG mRNA sequence features-based random forest classifier for prediction of transcription rate fold-changes in the mouse *Mecp2 y/-*. H, 3’UTR or CDS mRNA sequence features-based random forest classifier for prediction of transcription rate fold-changes in the mouse RTT model. I, ROC and PRC showing overall performance of the classifier for prediction of mouse transcription rate fold-change based on the nuclear dataset. TR= transcription rate, HL= half-life, SS= steady-state.

**Figure S3.**
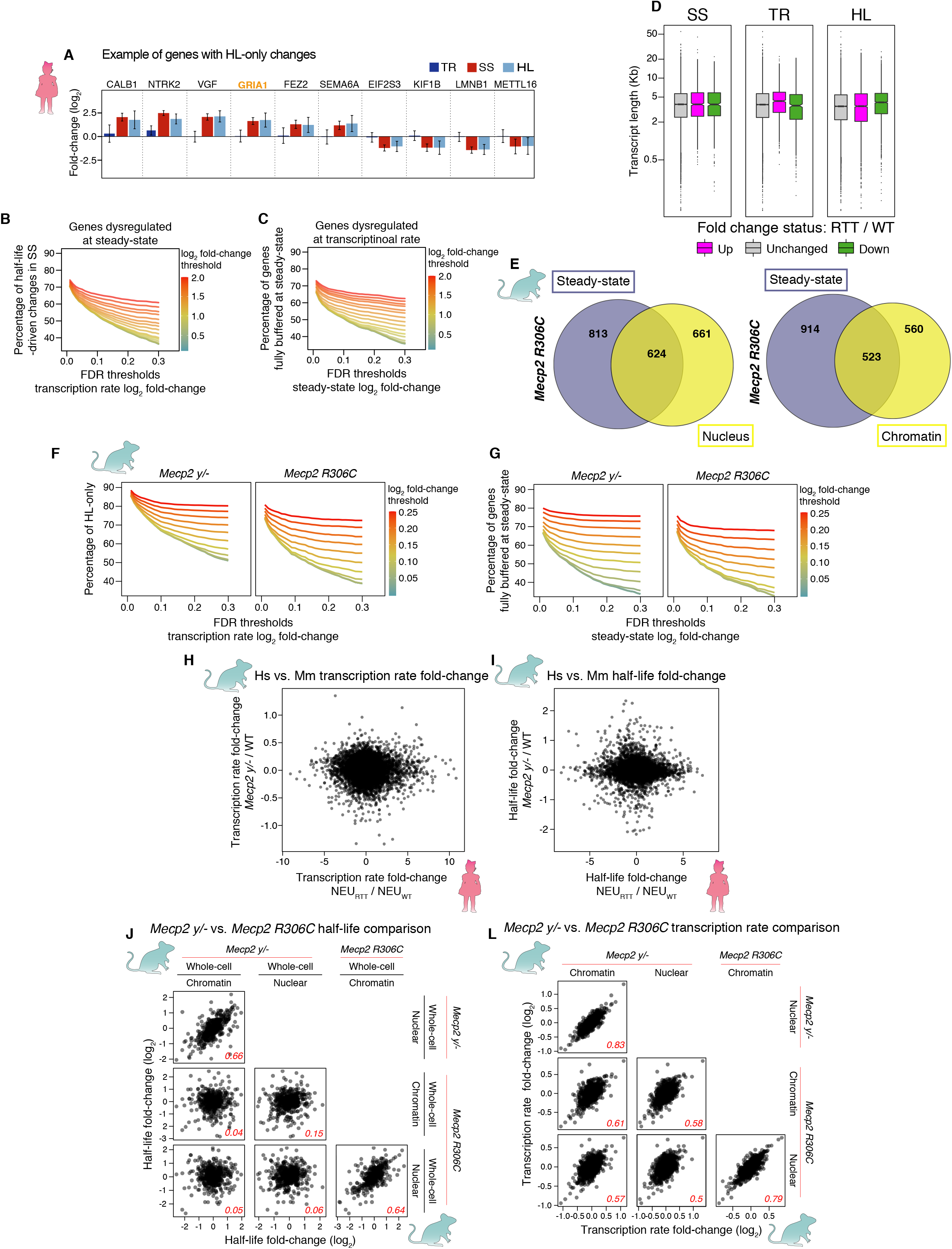
Human and mouse RTT models show half-life changes in steady-state mRNA levels. A, example of genes with mRNA half-life only changes leading to changes in the steady-state independent of transcription rate. GRIA1 is known to confer a high risk for autism when mutated. B, percentage of genes with no significant transcription rate shift or HL-only changes in steady-state (y-axis) as a function of the FDR (x-axis) and fold-change (color) thresholds. C, percentage of genes with no significant steadystate shift or fully buffered by mRNA stability mechanisms (y-axis) as a function of the FDR (x-axis) and fold-change (color) thresholds. D, human genes with steady-state, transcription rate, or half-life foldchanges in NEU_RTT_ do not show significant enrichment for transcript length as seen for adult mice^9^. E, the overlap of genes altered at transcription rate based on nuclear (left panel) or chromatin-associated RNA (right panel) and steady-state in the *Mecp2 R306C* mouse neurons. F, percentage of genes with no significant steady-state shift or fully buffered by half-life (y-axis) as a function of the FDR (x-axis) and fold-change (color) thresholds in the *Mecp2* y/- and *Mecp2 R306C* mice. G, percentage of genes with no significant transcription rate shift or with half-life change (y-axis) as a function of the FDR (x-axis) and fold-change (color) thresholds in the *Mecp2* y/- and *Mecp2 R306C* mice. H-I comparison of the foldchanges between genes in human (Hs) and mouse (Mm) RTT models showing limited agreement in the identity of genes with altered transcription rate (H) and half-life (I) between species. J-L, the identity of genes differentially regulated at half-life and transcription rate is also limited when comparing the *Mecp2* y/- and *Mecp2 R306C* mouse models. R values for each comparison are depicted inside boxes in red fonts. TR= transcription rate, HL= half-life, SS= steady-state.

**Figure S4.**
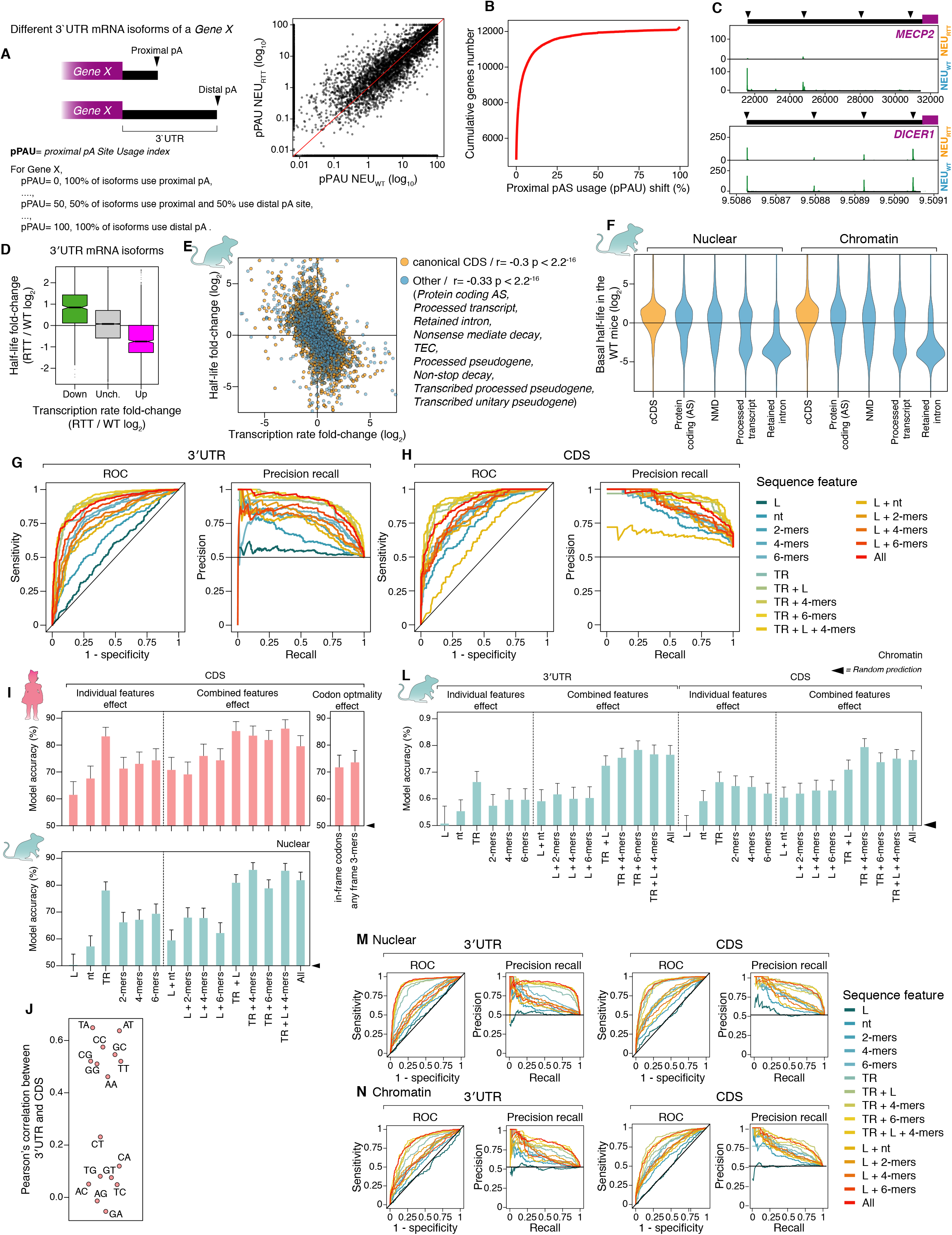
Quantification of 3’UTR and alternatively-spliced isoforms in humans and mice, and predictive models of half-life changes in mice. A, Overall schematics for quantification of alternative poly-Adenylation (APA) changes between NEU_WT_ and NEU_RTT_. Proximal poly-Adenylation site usage index (pPAU) quantifies the percentage of mRNA isoforms cleaved and poly-Adenylated at the proximal pA site, where 0 or 100 means that all mRNA isoforms of a specific gene use the proximal site (short 3’UTR isoform) or the distal (long 3’UTR isoform), respectively. Scatter-plot (right panel) shows a high correlation between pPAU values between human NEU_WT_ and NEU_RTT_. B, cumulative probability plot showing that the vast majority of genes show small changes <10% in pPAU index between human NEU_WT_ and NEU_RTT_. C, representative sequencing read peaks showing absence of *MECP2* 3’UTR reads in the NEU_RTT_ sample (upper) and peaks corresponding to *DICER1* polyadenylation sites (arrowheads) in the 3’UTR as an example of genes with similar pPAU index values in NEU_WT_ and NEU_RTT_ samples. Y-axis represents number of sequencing reads. D, quantification of mRNA at the 3’UTR isoform level indicating that most expressed 3’UTR isoforms also undergo half-life buffering. E, analysis of the alternatively-spliced mRNA isoforms in the *Mecp2* y/- mouse model shows that all canonical CDS and non-protein-coding isoforms also undergo half-life buffering, indicating that changes in mRNA splicing are not the underlying mechanism regulating the changes in mRNA half-life in response to transcription rate changes. F, quantification of the basal halflife of mRNA isoforms in the WT mice, as measured using either nuclear (left) or chromatin-associated RNA (right), confirms that non-sense mediated decay (NMD), processed transcripts, and retained intron mRNA isoforms are in general less stable (lower half-life) than the canonical CDS (cCDS) and alternatively-spliced (AS) protein-coding isoforms. This underscores the quality of the mRNA half-life calculations based on the ratios of whole-cell versus nuclear or chromatin-associated mRNAs. G and H, ROC and PRC graphs showing overall performance of the classifier for prediction of half-life fold-changes based on 3’UTR (G) or CDS (H) regions of the mRNAs. I, CDS specific mRNA-sequence features based random forest classifier for prediction of half-life fold-change for the human and *Mecp2* y/- mouse model. J, Pearson’s correlation between CDS and 3’UTR sequence of buffered genes indicating significant similarities between the sequence composition in these two regions. L, mRNA-sequence features based random forest classifier for prediction of half-life fold-change based on the chromatin-associated RNA in the *Mecp2* y/- mouse model. Similar to the half-life fold-change based on the nuclear-associated RNA, both 3’UTR and CDS contain sequence elements that can predict half-life changes with high accuracy. M and N, ROC and PRC graphs showing overall performance of the classifier for prediction of half-life fold-change in the *Mecp2* y/- mouse model based on nuclear (M) or chromatin-associated RNA (N). TR= transcription rate, HL= half-life, SS= steady-state

## Methods

### iPSC cultures and neuronal differentiation

iPSC lines #37 (WT) and #20 (isogenic *MECP2*-null) from a female patient were previously described^21^. Both cell lines were generated and cultured under the approval of the SickKids Research Ethics Board and the Canadian Institutes of Health Research Stem Cell Oversight Committee. iPSC lines were cultured in 5% CO2 on BD hESC-qualified matrigel (BD) in mTeSR medium (STEMCELL Technologies). Cultures were passaged using ReLeSR (STEMCELL Technologies) following the manufacture’s instruction every 6-7 days. For neuronal induction, iPSCs were aggregated as Embryoid Bodies (EBs) in low-attachment dishes in N2 media containing laminin (1 ml/ml) with 10 mM SB431542, 2 mM DSM, and 1x penicillin-streptomycin changed daily. After 7 days, EBs were plated on poly-L-ornithine + laminin-coated dishes and grown in N2 media + laminin (1 ml/ml). After 7 days, neural rosettes were manually picked and transferred to poly-L-ornithine + laminin-coated wells. After 7 days, neural rosettes were picked a second time, digested with Accutase and plated on poly-L-ornithine + laminin-coated wells. Resulting neural precursor cells (NPC) were grown as a monolayer and split every 5-7 days in NPC media containing DMEM/F12, N2, B27, 1x non-essential amino acid (NEAA), 2 mg/ml Heparin, 1 mg/ml laminin. To generate neurons, NPCs were plated on poly-L-ornithine + laminin-coated plates at a density of 10^6^ cells per 10 cm dish and cultured for 3 weeks in neural differentiation medium (Neurobasal, N2, B27, 1 mg/ml laminin, 1x penicillin-streptomycin, 10ng/ml BDNF, 10ng/ml GDNF, 200 mM ascorbic acid, and 10 mM cAMP).

### Neuronal enrichment using MACS

Neuronal cultures were enriched for all experiments to exclude contaminating glia and neuro progenitor cells present after differentiation. Enrichment of 3-week old neuronal cultures was made as described earlier^33,34^. 3-week old heterogeneous neuronal cultures were enriched by a negative selection strategy using antibodies against surface markers CD44 and CD184 (recognizing NPCs, glial progenitors and astrocytes)^35^ using magnetic-activated cell sorting (MACS^®^ - Miltenyi Biotec). After enrichment, neurons were re-seeded onto Matrigel-coated 6-well plates, cultured in neural differentiation medium, and allowed to recover for one extra week, for a total of 4 weeks neuronal differentiation.

### 4sU metabolic labeling of Neurons and RNA extractions

When enriched neuronal cultures reached 4-week of differentiation, media was replaced with neuronal differentiation media supplemented with 100 μM 4sU (Sigma-Aldrich) reconstituted in DMSO. Neurons were harvested at 0.5, 1, 4, 8, and 24 h after the addition of 4sU (except for the *MECP2*-null line where the time-point 1h was omitted from both replicates due to low differentiation yields). Metabolic labeling was designed such that all time points were collected together. After incorporation, cells were quickly washed twice with ice-cold 1× PBS Total RNA and scraped into ice-cold 1.5 ml Eppendorf tubes. Cells were collected by spinning at 1000g for 5 min at 4°C and cell pellets were resuspended in 1 mL of Trizol (Thermo Fisher Scientific). Total RNA was extracted according to manufacturer instructions. The SS sample was prepared from a 5μg aliquot of the 24 h time-point added with 0.5 μg of both 4sU labeled and unlabeled spike-in RNAs. Neuronal viability in the presence of 100 μM 4sU was monitored up to 24 h of treatment on parallel cultures by using Trypan blue staining and live/dead cell counting.

### Biotinylation and pull down of 4sU-labeled RNAs

50 μg of total neuronal RNA was mixed with 5 μg unlabeled *yeast* RNA and 5 μg 4sU-labeled S2 *fly* RNA in a total volume of 120 μL. 1 mg/mL HPDP-biotin (ThermoFisher Scientific) was reconstituted in dimethylformamide by shaking at 37°C for 30 min at 300 RPM. 120 μL of 2.5× citrate buffer (25 mM citrate, pH 4.5, 2.5 mL EDTA) and 60 μL of 1 mg/mL HPDP-biotin were added to the premixed RNA sample for each time point. The solution was incubated at 37°C for 2 h at 300 RPM on an Eppendorf ThermoMixer F1.5 in the dark. Samples were extracted twice with acid phenol, pH 4.5, and once with chloroform. RNA was precipitated with 18 μL 5M NaCl, 750 μL 100% ethanol, and 2 μL GlycoBlue (Invitrogen) overnight at −20°C. Precipitated RNA was pelleted for 30 min at 21,000*g* at 4°C. The RNA pellet was resuspended in 200 μL of 1× wash buffer (10 mM Tris-HCl, pH 7.4, 50 mM NaCl, 1 mM EDTA). Biotinylated RNA was purified using the μMACS Streptavidin microbeads system (Miltenyi Biotec). 50 μL Miltenyi beads per sample were pre-blocked with 48 μL 1× wash buffer and 2 μL yeast tRNA (Invitrogen), rotating for 20 min at room temperature. μMACS microcolumns were washed 1× with 100 μL nucleic acid equilibration buffer (Miltenyi Biotec), followed by 5× washes with 100 μL 1× wash buffer. Beads were applied to microcolumns in 100 μL aliquots and again washed 5× with 100 μL 1× wash buffer. Beads were demagnetized and eluted off the column with 2× 100 μL 1× wash buffer, and columns were placed back on the magnetic stand. A total of 200 μL beads was mixed with each sample of biotinylated RNA and rotated at room temperature for 20 min. Samples were applied to the microcolumns in 100 μL aliquots, washed 3× with 400 μL wash A buffer (10 mM Tris-HCl, pH 7.4, 6 M urea, 10 mM EDTA) prewarmed to 65°C, and washed 3× with 400 μL wash B buffer (10 mM Tris-HCl, pH 7.4, 1 M NaCl, 10 mM EDTA). RNA was eluted with 5× 100 μL of 1× wash buffer supplemented with 0.1 M DTT, and flow-through was collected in a tube. Purified RNA was precipitated with 30 μL 5 M NaCl, 2 μL GlycoBlue, and 1 mL 100% ethanol, incubated at −20°C overnight. Samples were spun at 21,000*g* at 4°C for 30 min and resuspended in 20 μL water. RNA quality was assessed by running 3 μL of samples on a 1.5% agarose gel.

### Transcription rate measurement using EU

Transcription rate measurements were validated by an alternative method using the metabolic incorporation of 5-ethynyl uridine (5-EU) followed by quantifying mRNA levels by qRT-PCR. NEU_WT_ and NEU_RTT_ were incubated with 0.5mM 5-EU (ThermoFisher) for 30 min. Total RNA was extracted and processed using Click-iT Nascent RNA Capture Kit (ThermoFisher) according to the manufacturer’s instructions. The captured RNAs were used as a template for cDNA synthesis followed by qRT-PCR to quantify mRNA level (see primer list below). Genes were chosen to cover a wide range of transcription rate changes determined by RATE-seq.

### Half-life measurements using transcription blocking

Half-life measurements were validated by an alternative method using transcription blocking followed by quantifying mRNA levels by qRT-PCR. 10μg/mL actinomycin D (Sigma-Aldrich) was added to NEU_WT_ and NEU_RTT_. RNAs were isolated at 1h, 3h, and 6h time points using the RNeasy Plus kit (QIAGEN). The RNAs were used as a template for cDNA synthesis followed by qRT-PCR to quantify mRNA level. Genes were chosen to cover a wide range of half-life changes as determined by RATE-seq.

### cDNA synthesis and qRT-PCR

cDNAs were synthesized using SuperScript III reverse transcriptase (ThermoFisher) with random hexamer primers according to the manufacturer’s instructions. For qRT-PCR, we used SYBR Select PCR Master Mix (ThermoFisher). Fold-changes were calculated by the ΔΔCt methods using Glyceraldehyde 3-phosphate dehydrogenase (GAPDH) and 18S as housekeeping genes, averaged between technical and subsequently biological replicates to achieve an average fold difference.

### miRNA extraction and spike-in strategy

To calculate relative and absolute differences in the miRNA population in NEU_WT_ and NEU_RTT_, small RNAs were extracted from two replicates of both lines using the same number of cells followed by the addition of a set of spike-in RNAs. Small RNAs were extracted from 500,000 neurons of each line using the SPLIT RNA extraction Kit (Lexogen) according to the manufacturer’s instructions. A set of 52 RNA spike-ins (QIAseq miRNA Library QC Spike-Ins – Qiagen) that spanned a wide range of concentrations were added to the recovered RNAs according to the manufacturer’s instructions. Sequencing libraries were made using the Small RNA library preparation kit NEBNext (NEB) according to the manufacturer’s instructions. Sequencing was performed on the Illumina HiSeq 2500 using the Rapid Run mode. Datasets are available upon request

### Library preparation and RNA-sequencing

RNA-seq libraries were prepared for each time-points and seady-state sample using the QuantSeq 3’ mRNA-Seq Library Prep Kit FWD for Illumina (Lexogen) automated on the NGS WorkStation (Agilent) at The Centre for Applied Genomics (TCAG) according to the manufacturer’s instructions. PCR cycle numbers were determined using the PCR Add-on Kit for Illumina (Lexogen). All steady-state samples were processed with 250 ng of total RNA input. To minimize variability between time-points within a batch, RNA samples were processed with the same total RNA input with a minimum of 100 ng of total RNA used. Each sample was spiked-in with ERCC RNA Spike-In Control Mix 1 (Ambion) according to the manufacturer’s instructions prior to the start of library preparation. Library quality and quantity were measured at TCAG prior to sequencing with Bioanalyzer (Agilent) and KAPA qPCR (Roche). Sequencing was also performed at TCAG on the Illumina HiSeq 2500 with single-end 100bp read length yielding 40 to 50 million reads. Datasets are available upon request.

### Processing of raw sequencing reads

Processing starts with trimming of reads in 4 steps using cutadapt version 1.10^36^. First, we removed adapters exactly at the 3’-end of the reads (-a AGATCGGAAGAGCACACGTCTGAACTCCAGTCAX -O 4 -e 0.1 --minimum-length 25). Second, we removed internal or long stretches of adapter (-a AGATCGGAAGAGCACACGTCTGAACTCCAGTCA -O 30 -e 0.18 --minimum-length 25). Third, we trimmed low-quality bases at the 3’-end of the reads (-q 20 -O 4). Finally, we removed poly-A tail at the 3’-end of the reads (-a AAAAAAAAAAAAAAAAAAAAAAAAAAAAAAX -O 4 -e 0 –minimumlength 25).

### Generation of custom hybrid genome index and reads alignment with STAR

We generated a custom genome index to accommodate quantification of yeast, fly, and ERCC spike-in RNA. Annotations (gencode version 29, flybase version all-r6.22, saccharomyces_cerevisiae.gff from yeastgenome.org, custom for ERCC) and genomes (hg38, dm6, sacCer3, ERCC from ThermoFisher) for all species and ERCC were combined and then processed with STAR version 2.6.0c (--sjdbOverhang 100). Finally, reads are aligned to hybrid genome with STAR version 2.6.0c (default settings)^37^.

### Quantification of RNA abundance

Poly-A sites were obtained from PolyA_DB version 3 and converted to hg38 coordinates with *liftOver* (UCSC)^38,39^. Reads with MAPQ < 2 are filtered out. Finally, usage of poly-A sites was defined as a sum of reads whose 3’-ends are falling within 20bp upstream and 10bp downstream of the poly-A sites. The sum was counted with a custom Python script using *pybedtools, pysam, pypiper*^40–42^. Annotation of pri-miRNA transcripts structures was downloaded from Mendel lab^43^. Each transcript was matched to miRNA gene based on overlaps with GENCODE annotated pre-miRNA coordinates^44^. Then, pri-miRNA poly-A sites overlapping mRNA or lncRNA poly-A sites from PolyA_DB were removed. Finally, usage of pri-miRNA poly-A sites was quantified with *featureCounts* (strandSpecific=1, read2pos=3 from *Rsubread* package)^45^. The abundance of mature miRNAs was quantified with *mirdeep2* pipeline^46^. Reads were preprocessed and collapsed with *mapper.pl* script (-e -h -j -k AGATCGGAAGAGCACA -l 18 -m -v) and quantified with *quantifier.pl* script, using hairpin and mature sequences obtained from miRbase^47^.

### Normalization of read counts

Human raw counts for each sample are divided by the total abundance of fly spike-in RNA, estimated as a sum of primary alignments to the fly genome. This normalization reconstructs the fraction of labeled human RNA at each timepoint.

### Transcription rate and half-life measurements

The transcription rate was estimated from early time points. First, normalized counts at 1 hour were divided by 2 to create a new approximate replicate at 30 mins timepoint. This assumes that RNA degradation is negligible for most genes at early time points. Then, before comparing cell types with a *DESeq2*, counts are further quantile normalized between replicates of the same cell type with *normalize.quantiles (preprocessCore* package)^48,49^. Half-life was estimated in 2 separate ways: fit of a 4sU saturation curve and the ratio of steady-state to transcription rate. For the 4sU saturation curve method, the half-life is estimated in a 2-step procedure. First, normalized counts are fit with *nls* (nonlinear least squares from *stats* package) to approximate the true number of counts Y at each timepoint. Then, in a second pass, normalized counts are fit again with *nls*, but now correcting for the increase in variance using weights set as 1/Y. Confidence intervals are estimated with *confint* function (*stats* package). For the ratio method, the half-life is estimated with *DESeq2* using raw human counts from 30 mins, 1 hour and steady-state timepoints (design =~ assay). In the assay factor, 30 mins and 1-hour samples correspond to the transcription rate.

### Processing of mouse datasets

Mouse data for whole-cell, nuclear and chromatin RNA-seq was downloaded from GSE128178. Mouse *Mecp2* ChIP-seq was downloaded from GSE139509. Differential expression analysis for nuclear and chromatin RNA-seq was downloaded from the supplementary materials of the Boxer *et al* study^9^. The abundance of 3’UTR isoforms for all samples is estimated using the QAPA standard pipeline^50^. Half-life was estimated as a ratio between whole-cell counts and nuclear or chromatin counts using the interaction term approach in *DESeq2* (design =~ celltype + batch + assay + celltype:assay). Coefficient of the celltype:assay term is used to measure the log2 fold-change in half-life between cell types. Before the *DESeq2* run, we filter out genes with a sum of counts in replicates less than 20 in a pair of compared cell types.

### Random forest prediction of up and down-regulated genes in transcription rates and mRNA half-life

Fold-changes in human half-life in log scale log_2_FC_HL_ between cell types A and B were estimated as follows:

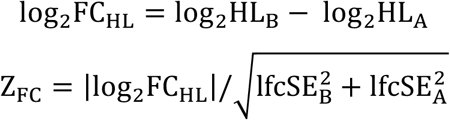

There, log_2_HL_A_ and lfcSE_A_ were average and standard error of the half-life in cell type A, estimated by *DESeq2* as a ratio from steady-state and transcription rate replicates. Z-score of a fold-change Z_FC_ was used as a measure of accuracy.

Features of the classifier are frequencies of k-mers in 3’UTR, coding sequence or gene-body, calculated using *oligonucleotideFrequency* (*Biostrings* package)^51^. In addition, the methylation status of CA and CG in the gene-body was added for mouse analysis from GSE139509. Predicted variable denotes genes that are either up or down-regulated in transcription rate or half-life. Thresholds for the human data were:

1. TR_up_: log_2_FC_TR_ > 1 & padj < 0.1
2. TR_down_: log_2_FC_TR_ < −1 & padj < 0.1
3. HL_up_: log_2_FC_HL_ > 1 & Z_FC_ > median(Z_FC_)
4. HL_down_: log_2_FC_HL_ < −1 & Z_FC_ > median(Z_FC_)

The half-life for the mouse was either from nuclear or chromatin. Thresholds for the mouse data were:

1. TR_up_: log_2_FC_TR_ > 0.1 & FDR < 0.1
2. TR_down_: log_2_FC_TR_ < −0.1 & FDR < 0.1
3. TR_not_: |log_2_FC_TR_| < 0.1 or FDR > 0.1
4. HL_up_: log_2_FC_HL_ > 1 & pvalue < quantile(pvalue, 0.2)
5. HL_down_: log_2_FC_HL_ < −1 & pvalue < quantile(pvalue, 0.2)

Data was split into 75% and 25% for training and test sets. The classifier is trained using *randomForest* (*randomForest* package)^52^. Precision-recall and receiver operator characteristic curves were obtained with *evalmod* (*precrec* package)^53^.

### Transite analysis of miRNAs and RBPs

Genes were split into up- or down-regulated according to their transcription rate and half-life foldchange. Then, we performed multiple comparisons of 3’UTR sequences between groups of genes using *run_kmer_tsma (transite* package)^54^. Groups of compared genes:

1. Foreground: TR_down_ and HL_up_ Background: TR_down_
2. Foreground: TR_up_ and HL_down_ Background: TR_up_
3. Foreground: TR_not_ and HL_down_ Background: TR_not_

These comparisons were performed for all transite RNA binding protein motifs and for TargetScan seed sequences^24^. TargetScan analysis includes 439 human miRNA-seed sequences with family conservation scores of 0,1,2 that were selected from miR_Family_Info.txt (TargetScan website). Definitions of up- and down-regulated genes:

6. TR_up_: log_2_FC_TR_ > 0.5 & padj < 0.1
7. TR_down_: log_2_FC_TR_ < −0.5 & padj < 0.1
8. TR_not_: |log_2_FC_TR_| < 0.5 or padj > 0.1
9. HL_up_: log_2_FC_HL_ > 1 & Z_FC_ > median(Z_FC_)
10. HL_down_: log_2_FC_HL_ < −1 & Z_FC_ > median(Z_FC_)

The results of *transite* analysis were further processed with a custom script for multiple hypothesis correction. Motifs with a low number of sites detected in both background and foreground were removed from the analysis. A separate threshold for the number of sites was chosen for each transite analysis. A threshold was determined from a requirement for p-values distribution to be unimodal and enriching at p=0. The distribution of p-values with unfiltered sites is bimodal with peaks at both p=0 and p=1.

### Primer list for qRT-PCRs

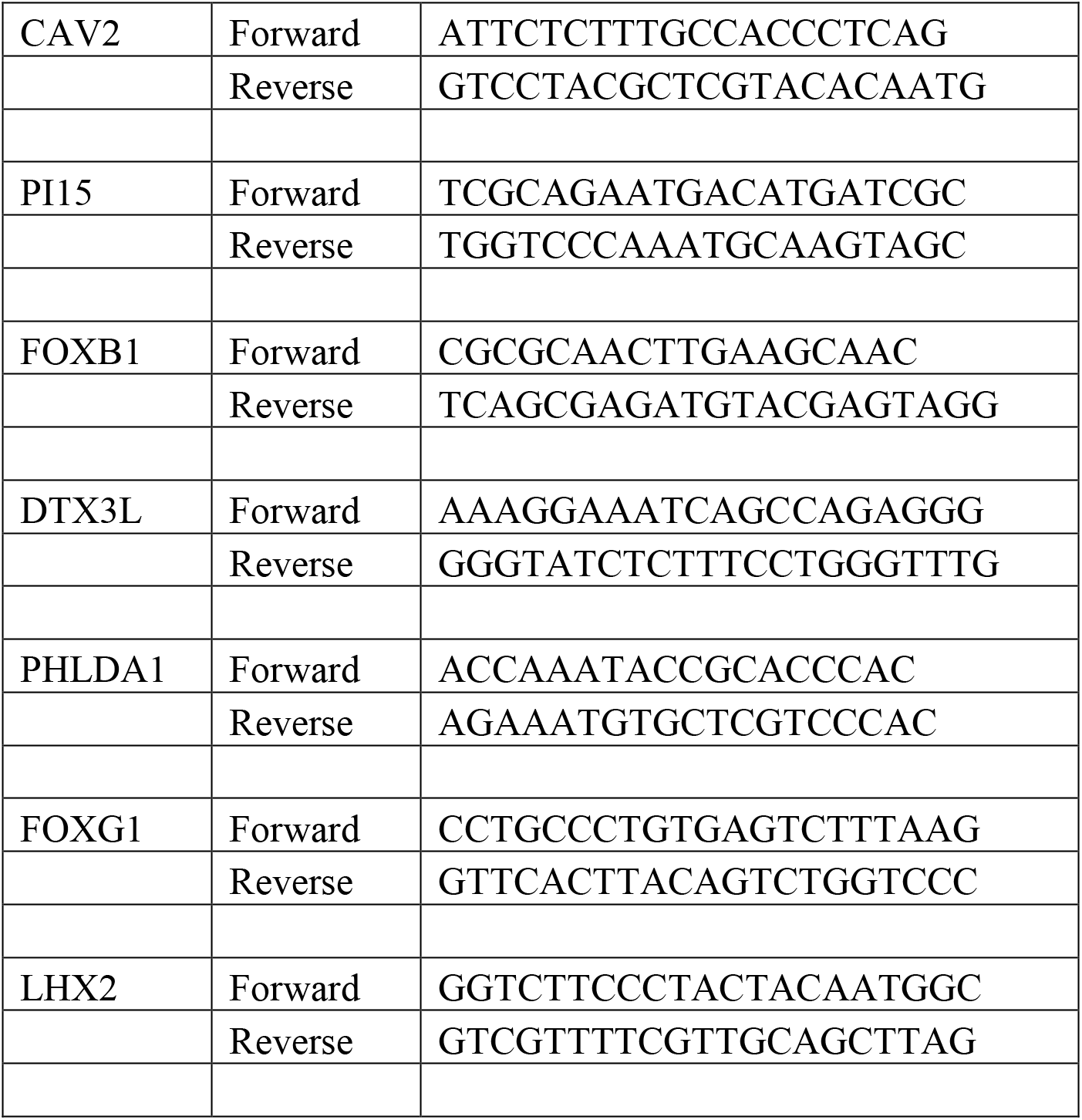
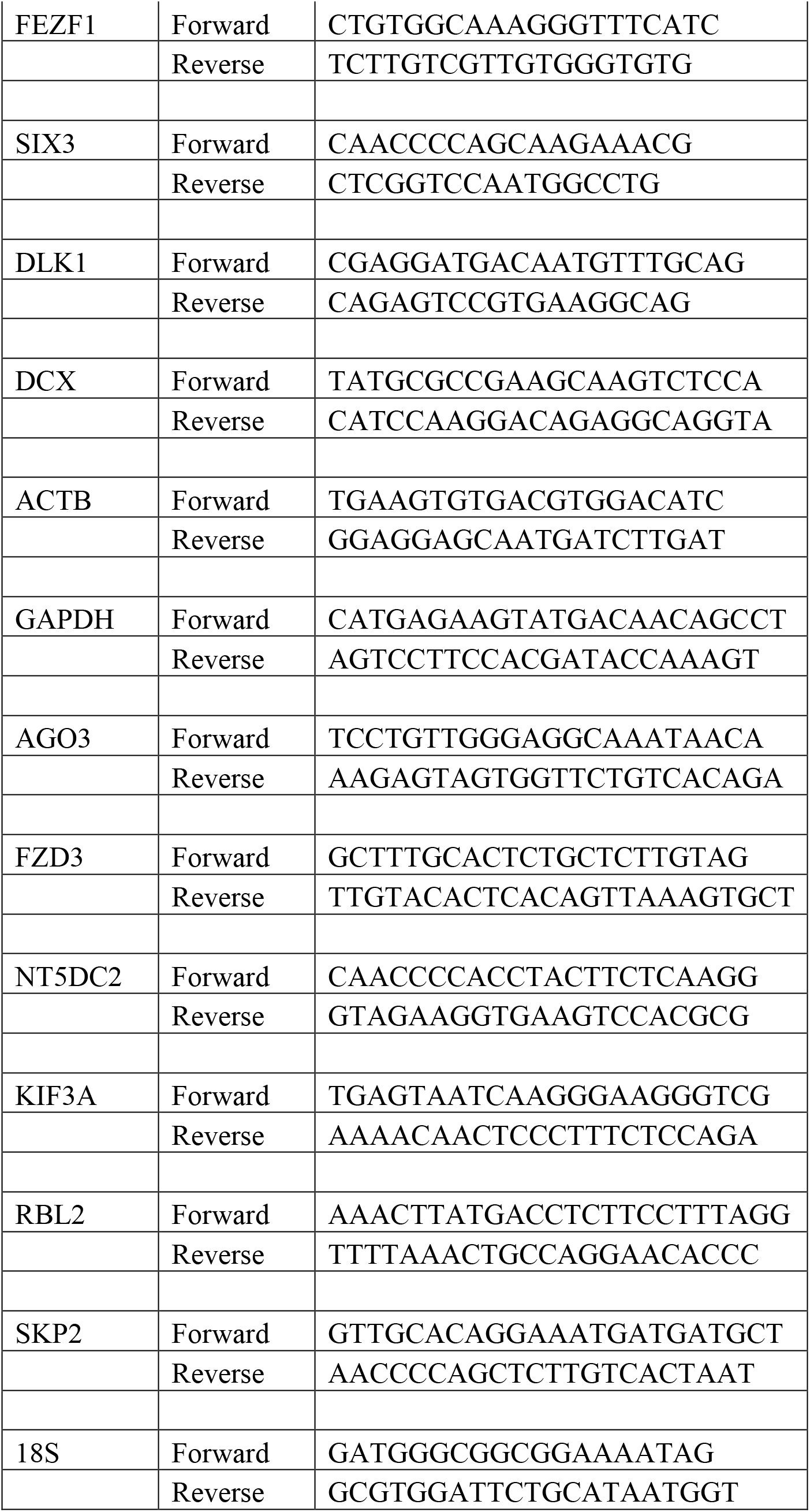

